# Critical role of CD206+ macrophages in promoting a cDC1-NK-CD8 T cell anti-tumor immune axis

**DOI:** 10.1101/2023.10.31.560822

**Authors:** Arja Ray, Kenneth H. Hu, Kelly Kersten, Tristan Courau, Nicholas F. Kuhn, Itzia Zaleta-Linares, Bushra Samad, Alexis J. Combes, Matthew F. Krummel

**Affiliations:** Department of Pathology, University of California, San Francisco, CA 94143, USA; ImmunoX Initiative, University of California, San Francisco, CA 94143, USA; UCSF CoLabs, University of California, San Francisco, CA 94143, USA; Department of Medicine, University of California, San Francisco, CA 94143, USA

## Abstract

Tumor-associated macrophages (TAMs) are frequently categorized as being ‘M1’ or ‘M2’ polarized, even as substantial data challenges this binary modeling of macrophage cell state. One molecule consistently referenced as a delineator of a putative immunosuppressive ‘M2’ state is the surface protein CD206. We thus made a novel conditional CD206 (*Mrc1*) knock-in mouse to specifically visualize and/or deplete CD206+ ‘M2-like’ TAMs and assess their correspondence with pro-tumoral immunity. Early, but not late depletion of CD206+ macrophages and monocytes (here, ‘Mono/Macs’) led to an indirect loss of a key anti-tumor network of NK cells, conventional type I dendritic cells (cDC1) and CD8 T cells. Among myeloid cells, we found that the CD206+ TAMs are the primary producers of CXCL9, and able to differentially attract activated CD8 T cells. In contrast, a population of stress-responsive TAMs (“Hypoxic” or *Spp1*+) and immature monocytes, which lack CD206 expression and become prominent following early depletion, expressed markedly diminished levels of CXCL9. Those NK and CD8 T cells which enter CD206- depleted tumors express vastly reduced levels of the corresponding receptor *Cxcr3,* the cDC1- attracting chemokine *Xcl1* and cDC1 growth factor *Flt3l* transcripts. Consistent with the loss of this critical network, early CD206+ TAM depletion decreased tumor control by antigen specific CD8 T cells in mice. Likewise, in humans, the CD206^Replete^, but not the CD206^Depleted^ Mono/Mac gene signature correlated robustly with CD8 T cell, NK cell and stimulatory cDC1 gene signatures and transcriptomic signatures skewed towards CD206^Replete^ Mono/Macs associated with better survival. Together, these findings negate the unqualified classification of CD206+ ‘M2-like’ macrophages as immunosuppressive by illuminating contexts for their role in organizing a critical tumor-reactive archetype of immunity.

## Introduction

Macrophages have diverse roles in homeostasis and disease and a refined understanding of the direct and indirect effects of targeting them in tumors is an imperative, given the current impetus in developing myeloid targeting therapies for cancer(*1, 2*). A widely used shortcut for describing macrophage function in tumors involves an ‘M1’ versus ‘M2’ nomenclature, derived from in vitro skewing with Th1 versus Th2 cytokines, and often equated with pro and anti-inflammatory functions respectively. However, this binary M1/M2 delineation of macrophage phenotype does not capture the heterogeneity at the single cell level (*3–5*). In addition, there is scant evidence of these markers being part of coordinated gene programs in vivo. In wound healing, *Arg1* and *Mrc1*(gene corresponding to the mannose-binding C-type lectin CD206), both purportedly key markers of an M2 state, have distinct expression patterns (*6*). In both mouse and human tumors, there is also a complete lack of correlation among genes characterizing M1 or M2 phenotypes within Mono/Macs (*3*). In fact, M1 and M2 signatures in Mono/Macs often show correlated instead of opposing expression patterns in tumors (*4, 5*). Nonetheless, myeloid cells expressing CD206, sometimes therefore designated as ‘M2-like’, continue to be used as a marker of an immunosuppressive state. The detrimental effects of tumor-associated macrophages (TAMs) on anti-tumor immunity have indeed been highlighted by a number of critical studies (*7–11*). However, a holistic dissection of the role of CD206-expressing Mono/Macs and the precise effects of targeting them in tumors in vivo is lacking. We therefore developed a conditional knock-in reporter mouse using the *Mrc1* (CD206) allele that allows specific visualization and depletion of those cells to define their true impact on anti-tumor immunity.

## Results

To highlight CD206 surface expression variation across Mono/Mac differentiation in tumors, we identified relevant subsets from previously published single cell transcriptomics in B16F10 tumors (**Fig. 1A**, (*3*)), and applied flow cytometry to gate on those populations in a related B78chOVA ((*11*); B78: an amelanotic clone of B16 to allow imaging of tumors, chOVA: mCherry and Ovalbumin) tumor model. CD206 was most prominently expressed by terminal VCAM-1^hi^IL-7Rα^lo^ C1q TAMs (*3*) followed by the VCAM-1^lo^IL-7Rα^hi^ stress-responsive population (associated with enriched glycolysis, increased *Arg1* and *Il7r* expression(*3, 12*) and possibly hypoxic(*13*)), where TAMs were defined as CD45^+^Lin(CD90.2, Ly6G, B220, NK1.1, Siglec-F)^-^Ly6C^-^F480^+^CD24^-^ (**Supplementary Fig. S1A**). In contrast, CD206 was expressed at low levels by other less differentiated VCAM^-^IL-7Ra^-^ (DN) TAMs (**Fig. 1B, C**). Among monocytes, defined as CD45^+^Lin(CD90.2, Ly6G, B220, NK1.1, Siglec-F)^-^Ly6C^+^, the MHCII+ subset was the prominent CD206 expressor as opposed to early (immature, MHCII-) monocytes, albeit at lower levels than Stress and C1q TAMs (**Fig. 1B, C**). Overall, this analysis showed that CD206 is variably expressed across multiple monocyte and macrophage subsets, and generally increases with differentiation. However, when considering the use of this protein and the gene encoding it as a means of eliminating these macrophages and thereby studying their function, we noted that this scavenger receptor is frequently also expressed on other cells including endothelial cells and keratinocytes (*14, 15*) and may further be ectopically produced by other cells in the TME.

**Fig. 1:**
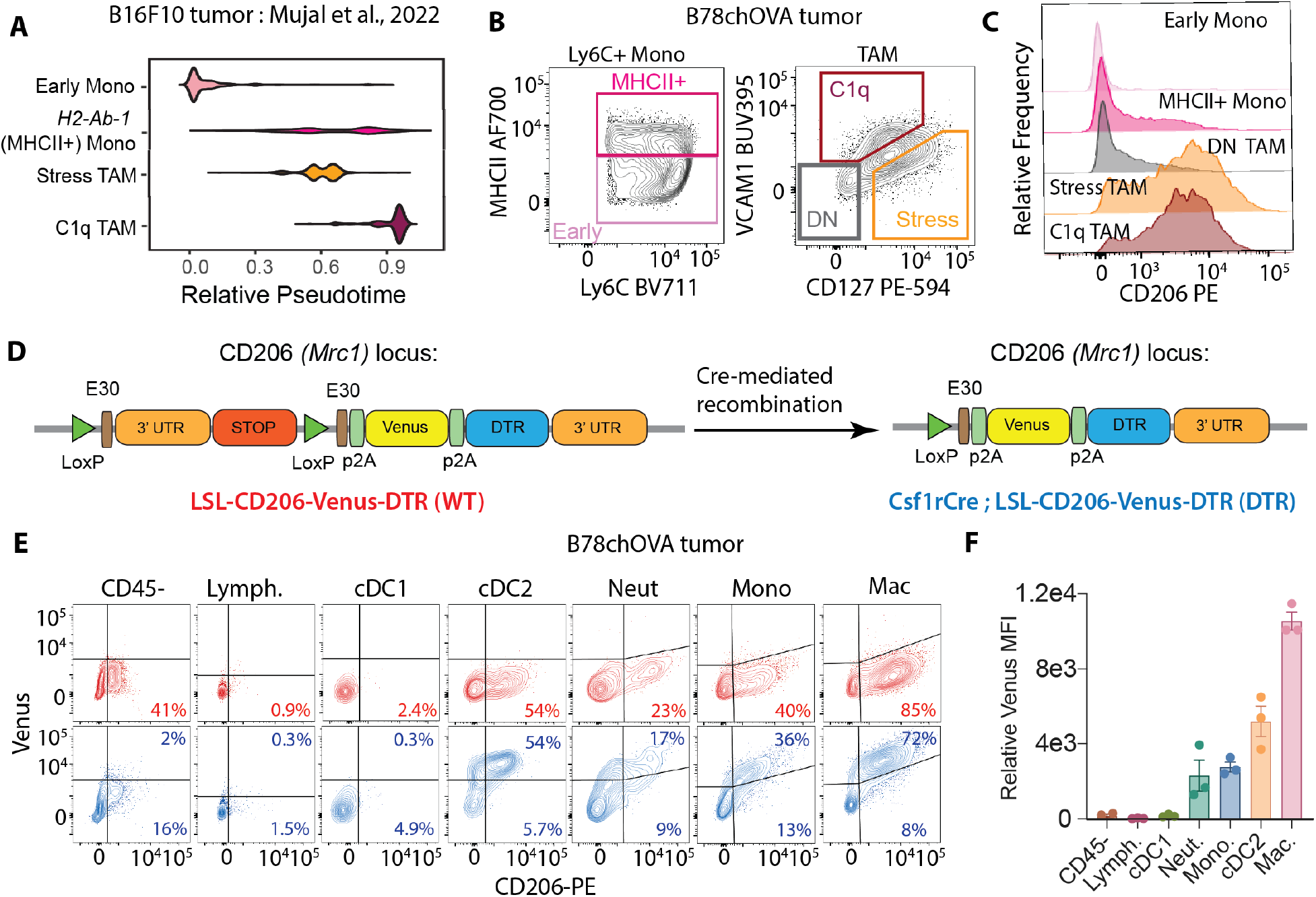
Genetic Myeloid-specific Labeling of CD206+ Macrophages in Tumors: **(A)** Pseudotime plots of select Mono/Mac subsets in B16F10 tumors from Mujal et al(*3*); **(B)** Gating on the equivalent subsets in B78chOVA tumors by flow cytometry and **(C)** CD206 expression in each of these subsets; **(D)** Schematic representation of the *Mrc1*^LSL-Venus-DTR^ knock-in construct before (WT) and after (DTR) Cre-mediated recombination by crossing to the *Csf1r*^Cre^ allele; **(E)** Flow cytometry plots showing reporter (Venus) and CD206 expression in different immune cells in d18 B78chOVA tumors in WT (red) and DTR (blue) mice with **(F)** quantification of relative reporter expression (DTR – WT) in the different subsets. data are mean +/− SEM, from 3 biological replicates, WT levels averaged from 2 biological replicates.

We thus generate a conditional system where a lineage-specific Cre could drive the recombination of a 3’ knock-in *Mrc1*^LSL-Venus-DTR^ allele (Venus : Yellow Fluorescent Protein variant for visualization; DTR: Diphtheria toxin receptor for depletion) (**Fig. 1D**). Then, using a *Csf1r*^Cre^; *Mrc1*^LSL-Venus-DTR^ cross (DTR), compared to a *Mrc1*^LSL-Venus-DTR^ (WT) control, we assessed the reporter expression in various immune and CD45- non-immune compartments in the subcutaneous melanoma model B78chOVA (**Fig. 1D, Supplementary Fig. S1A**). As predicted from CD206 expression, nearly 75% of TAMs showed robust Venus expression tightly correlated with surface expression of CD206 protein (**Fig. 1F**). We also found that nearly half of cDC2s, consistent with their monocytic origin as previously described(*16*), expressed Venus in this system. A small subset of CD206+ monocytes expressed the reporter, again consistent with *Csf1r* driven expression. Weak expression was also found in a yet smaller population of neutrophils. When viewed by the level of CD206-driven expression of the Venus marker, macrophages were 2-3x brighter than the other populations (**Fig. 1F**). Importantly, no reporter expression was detected in non-immune cells (which include endothelial cells and keratinocytes and where a small fraction is CD206+), lymphocytes and cDC1s (**Fig. 1E, F**). Likewise, in the proximal tumor-draining lymph nodes (tdLN), the same hierarchy of expression patterns was observed, albeit at much lower levels (**Supplementary Fig. S1B**). In addition, some tissue-resident macrophages, such as alveolar macrophages in the lung express high levels of CD206 and therefore the reporter, while interstitial macrophages, monocytes, neutrophils, and non-immune cells showed very modest to no expression (**Supplementary Fig. S1C, D**). Therefore, in *Csf1r*^Cre^; *Mrc1*^LSL-Venus-DTR^ mice, a robust marking of mature CD206+ macrophages in the subcutaneous tumor was observed, along with faithful marking of CD206+ subsets of monocytes, neutrophils and cDC2s, but not cDC1s, lymphocytes and non-immune cells.

### CD206+ TAM depletion leads to indirect loss of a reactive immune archetype

To test the impact of CD206+ macrophage targeting on the overall tumor immune microenvironment, we took advantage of the linkage of Venus and DTR expression in this background to deplete those cells. In our setup using subcutaneous B78chOVA tumors as described previously, adoptively transferred ovalbumin-specific OT-I cells allow the tracking of antigen-specific CD8 T cell responses, which nevertheless do not mediate tumor control(*11, 17*). We first confirmed that Cre-mediated induction of reporter expression without diphtheria toxin (DTx) administration did not alter the immune composition of these OVA-expressing tumors with OT-I transfer in the WT (*Mrc1*^LSL-Venus-DTR^) vs. DTR mice (*Csf1r*^Cre^; *Mrc1*^LSL-Venus-DTR^) (**Supplementary Fig. S2A**). With this baseline, we administered DTx either ‘late/acute’, namely in the last 4 days prior to the tumor harvest or ‘early/chronic’, i.e., every day 2-3 days, starting 2 days post T cell injection until harvest to parse out the role of CD206+ Mono/Macs in the TME (**Figure 2A**). These two modes of depletion represent perturbations at different phases of the establishment of the tumor immune microenvironment.

**Fig. 2:**
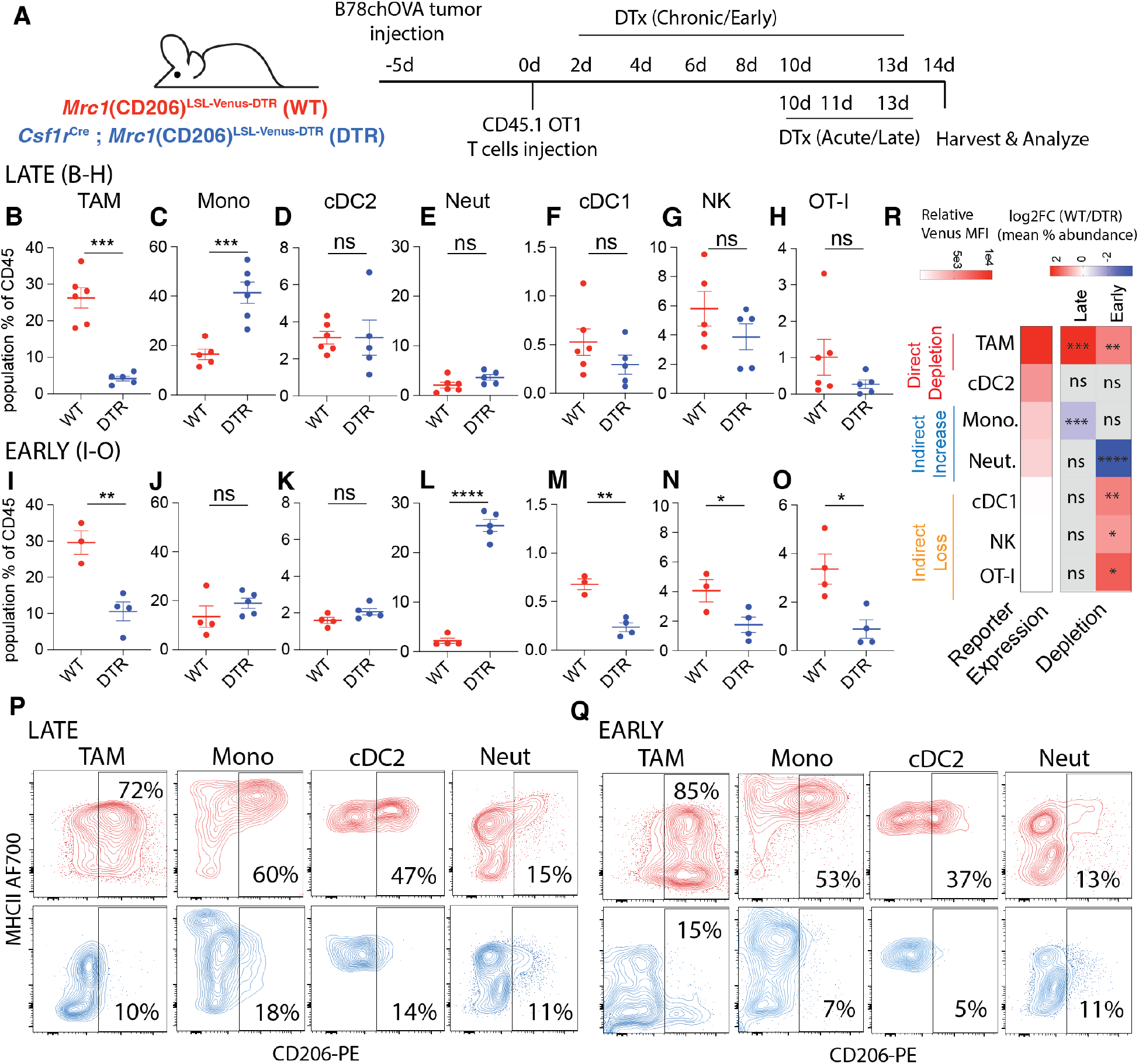
Early CD206+ TAM depletion leads to a coordinated and indirect loss of NK, cDC1 and CD8 T cells in the TME: **(A)** Schematic representation of the experimental setup for early and late CD206+ TAM depletion in B78chOVA tumors using *Mrc1*(CD206)^LSL-Venus-DTR^ (WT) and *Csf1r*^Cre^; CD206^LSL-Venus-DTR^ (DTR) mice; Relative abundance of different immune populations as a percentage of CD45+ cells with **(B-H)** late and **(I-O)** early depletion regimens; Representative flow cytometry plots showing CD206 vs. MHCII expression in different myeloid subsets in WT (red) and DTR (blue) mice in the (**P**) late and (**Q**) early depletion regimens. **(R)** heatmap representation of the log fold change of the ratio of mean abundances in WT and DTR mice (data from B-O), alongside the extent of reporter expression (mean relative Venus MFI from Fig. 1F) to indicate direct depletion and indirect loss or enrichment. (statistical significance is indicated on the respective squares; ***p <0.001, **p<0.01, *p <0.05, ns = no significance by Student’s t-tests).

In the context of late DTx administration, we found that this regimen specifically depleted the cells of interest and otherwise had little overall effect on other non-targeted cells. Thus, there was a strong reduction in the TAMs (**Figure 2B**) which corresponded to a specific loss of the CD206+ population (**Fig. 2P**). We found a compensatory rise in monocytes overall, accompanied by a specific loss of its CD206+ subset (**Figure 2C, 2P**). No significant loss was observed in cDC2s, cDC1s, NK cells, the adoptively transferred OTI, or neutrophils (**2D-H**), although the regimen did select against the CD206+ populations in the case of cDC2s and neutrophils (**Figure 2P**). Thus, direct depletion was reliably restricted to the highest expressors of the construct, namely the CD206+ TAM and monocyte populations.

When we depleted CD206+ populations with DTx early, i.e., starting 2d after OTI adoption, again we found robust depletion of TAMs (**Fig. 2I**) expectedly with a strong selection against those expressing CD206 (**Fig. 2Q**). As before, the CD206+ subsets of other populations were still depleted—robustly in monocytes and cDC2s and mildly in neutrophils (**Fig. 2K, Q**). A compensatory increase in neutrophils was observed, while overall abundance of monocytes, albeit biased towards CD206-, remained similar, in contrast to acute depletion. (**Fig. 2J, L**). Strikingly, under this early elimination regime, we also observed a decrease in intratumoral cDC1 abundance (**Fig. 2M**). Further, NK cells and transferred OT-I abundance in the tumor was also significantly compromised (**Fig. 2N, O**).

These trends in abundances were similar when expressed as percentage of live cells, indicating numerical changes, in all cases except the increase in monocytes following acute depletion (**Supplementary Fig. S2B-O**), which only trended higher. When we similarly treated non-tumor bearing DTR mice with DTx with six doses akin to the early depletion regimen in tumors, and analyzed the immune compositions in the skin (site of the ectopic tumor injections)(**Supplementary Fig. S2P**), no robust indirect loss of populations were observed, but an increase of neutrophils in an otherwise scarcely immune-populated skin was recorded (**Supplementary Fig. S2Q**). Given the associated increase in neutrophils, we repeated the same early depletion experiment in tumors, now with the addition of anti-Ly6G neutrophil depleting antibody or isotype control (**Supplementary Fig. S2R)** to assess whether the gained neutrophils played a role in the indirect loss of lymphocytes and cDC1s. As expected, both in terms of the total number of cells per gram of tumor (**Supplementary Fig. S2S)** and the percentage of CD45+ (immune) cells (**Supplementary Fig. S2T)**, the abundance of immune cell types in WT and DTR mice treated with isotype control mirrored those of the early depletion regimen. With anti-Ly6G treatment in DTR mice, neutrophils were reduced to levels below those of untreated controls, without any concomitant effect on the indirect depletion of cDC1s, NK cells and OT-I T cells (**Supplementary Fig. S2S, T**). Noting that none of these CD8, NK and cDC1 populations express the reporter, we concluded that a direct, targeted ablation of ‘M2-like’ CD206+ Mono/Macs by early DTx treatment in tumors led to the indirect loss of this key anti-tumor reactive archetype comprising of NK cells, cDC1s and antigen specific CD8 T cells(*18*), (**Fig. 2R**). This suggested that CD206+ Mono/Macs were involved in the recruitment and early establishment of this module in the TME.

### Early depletion skews Mono/Macs towards immature and hypoxic subsets

To define the macrophage subtypes associated with reactive immunity and their potential spatially segregated modes of action, we performed spatial transcriptomics of B78chOVA tumors, guided by Venus (CD206 reporter) expression. For this, we first spatially mapped the CD206+ macrophage population by Venus expression, using two-photon microscopy of B78chOVA tumor slices with transferred OT-I T cells marked by the CD2dsRed allele (**Fig. 3A**). Doing this revealed three distinct niches of CD206+ macrophage and T cell localization. The ‘edge’, which is macrophage and collagen-rich with modest T cell presence, ‘mid’, the interfacial layer with abundant T cell: macrophage interaction zones and ‘interior’. The interior is sparser in both immune cell types but represents the bulk of the tumor by volume (**Fig. 3A**). We then performed post-imaging spatial transcriptomics by ZipSeq (*19*) on CD45+ cells in these three zones (*11*) of B78chOVA tumors, with or without early DTx treatment harvested at d12 post T cell injection. UMAP projection of non-linear dimensional reduction and louvain clustering clearly showed the remarkable shift in tumor immune composition among control and DTx treated groups (**Fig. 3B, C, Supplementary Fig. S3A**). Notably, previously defined C1q and Stress-responsive (Stress) TAMs, which most robustly express CD206 at the protein level, along with MHCII+ and Interferon-stimulatory gene (ISG) -expressing monocytes were expectedly depleted by direct DTx action (**Fig. 3C**), with the robust depletion of C1q TAMs (VCAM-1^hi^IL-7Rα^lo^) and MHCII+ Monocytes verified by flow cytometry (**Fig. 3D**). On the other hand, early monocytes, neutrophils and a *Spp1*, *Hif1α*-expressing subset related to the Stress TAMs by shared expression of *Arg1, Il7r* (i.e., Stress^Spp1^ TAM), became prominent in the DTx treated condition (**Fig. 3C, Supplementary Fig. S3A**). The loss of cDC1:NK:CD8 populations was again evident in the analysis of relative abundance from the scRNASeq data (**Fig. 3C**). Even though the small area of the tumor edge was much denser in the CD206+ Mono/Macs as shown by imaging, the transcriptomic data suggests high CD206-expressing C1q and Stress TAMs were more or equally as abundant in the interior than the edge (**Fig. 3E**). consistent with the trajectory of increasing Mono/Mac differentiation towards the interior of the tumor(*11*). Overall, in this subcutaneous tumor model, the changes in immune subpopulations were not limited to a specific region of the tumor (**Fig. 3C**), but permeated throughout as a holistic overhaul of the tumor immune microenvironment.

**Fig. 3:**
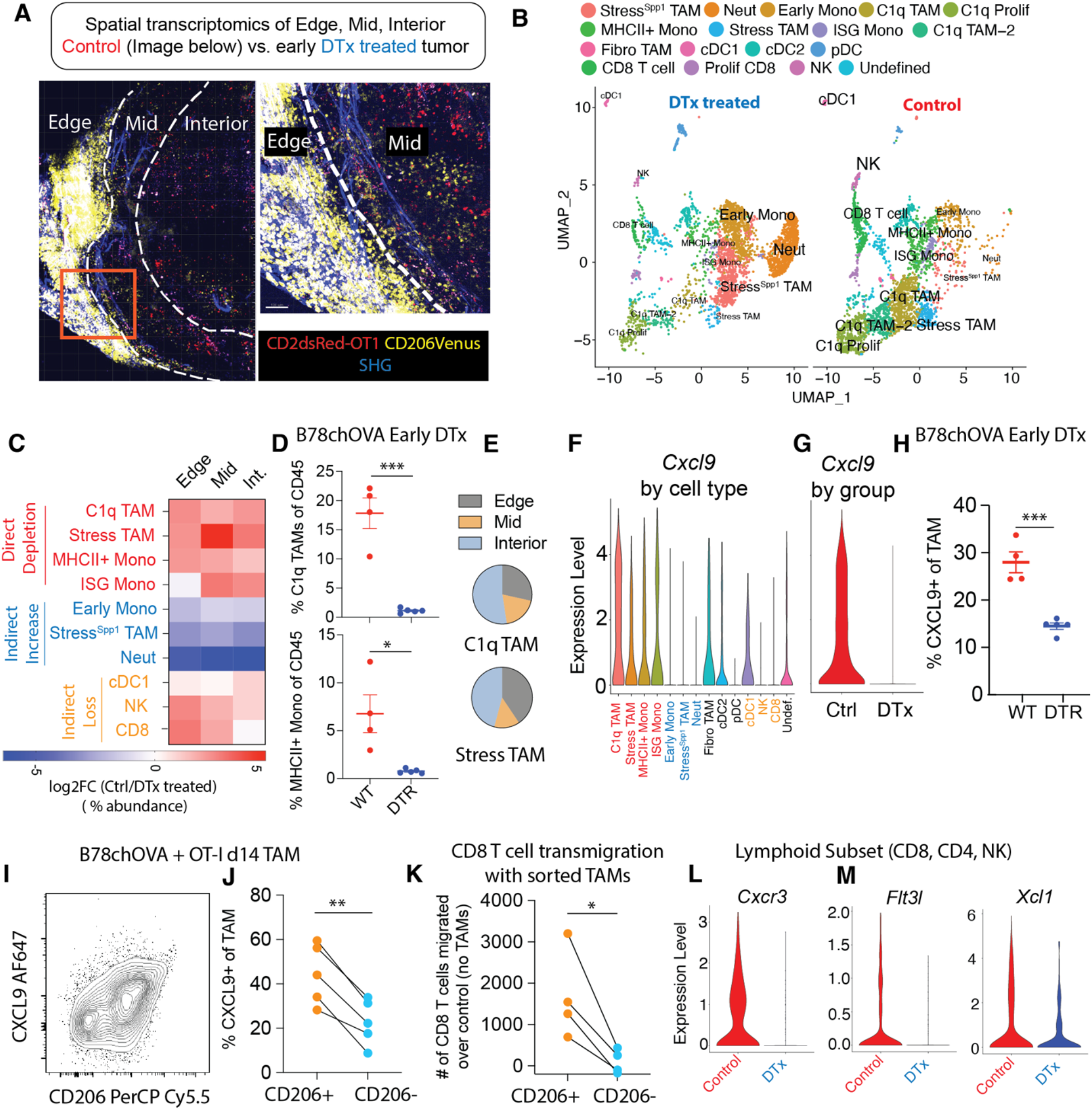
Loss of CXCL9-positive TAMs and CXCR3-expressing, cDC1 supportive lymphocytes with CD206+ TAM depletion. **(A)** Two-photon imaging of representative (control) B78chOVA tumors d12 post adoptive transfer of CD2dsRed; OT-I CD8 T cells showing three zones of Venus-expressing macrophage and associated CD8 T cell localization – edge, mid and interior (Int.) mapped by spatial transcriptomic barcoding ZipSeq; Boxed region is magnified (*right*) to show corresponding edge-mid interface SHG: Second Harmonic Generation; **(B)** UMAP representation of major immune cell populations obtained from Control and early DTx treated B78chOVA tumors d12 post OT-I injection aggregated across all three regions; **(C)** Summary heatmap showing relative log fold change of the abundance (calculated as the % of the total number of cells recovered within that region) of each major cluster in Ctrl/DTx treated conditions, split by region of tumor; *Cxcl9* expression; **(D)** Flow cytometry data showing abundance of C1q TAMs and MHCII+ Monocytes in Ctrl and DTx treated conditions; (**E**) Distribution of C1q and Stress-responsive TAMs in the three spatial regions in control B78chOVA tumors; *Cxcl9* expression **(F)** aggregated across treatment conditions by cluster and **(G)** aggregated across clusters by condition; **(I)** Representative flow cytometry plot showing intracellular CXCL9 vs. surface CD206 expression in TAMs in B78chOVA tumors at d14 post OT-I adoptive transfer and (**J**) the same CXCL9 expression split by CD206 positivity; **(K)** in vitro activated CD8 T cell migration at 3h through a 5µm transwell insert in the presence of sorted CD206+ vs. CD206- TAMs from B78chOVA tumors, normalized to migration with no TAMs; **(L)** *Cxcr3,* **(M)** *Flt3l* and *Xcl1* expression in the lymphocyte subset (CD8 T cell, NK cell and CD4 T cell) by treatment group. ***p <0.001, **p<0.01, *p <0.05, ns = no significance by Mann-Whitney test (D), unpaired t-test (H) and ratio paired t-test (J, K).

### CD206+ TAMs attract CXCR3-expressing, cDC1-supportive lymphocytes to the tumor

A well-established positive functional role of TAMs is the production of CXCL9 and CXCL10, inducing CXCR3-dependent lymphocyte recruitment in tumors(*20*). Given the indirect loss of lymphocytes upon early removal of CD206+ Mono/Macs, we hypothesized that this axis is prominent in CD206 positive myeloid populations. Analyzing the scSeq data in detail, we found that expression of *Cxcl10* (**Supplementary Fig. S3C**) and *Cxcl9* (**Fig. 3G**) in particular were markedly reduced in the DTx treated tumors and this corresponded to substantial expression by the directly depleted subsets (CD206+MHCII+ Mono/Macs) and none of the indirectly increased ones (Early Mono, Stress^Spp1^ TAM and Neutrophils) (**Supplementary Fig. S3B, Fig. 3F**). We confirmed this finding by flow cytometry for intracellular CXCL9 expression in TAMs from WT and DTR mice with early depletion (∼50% decline, **Fig. 3H, Supplementary Fig. S3D**). This analysis further revealed a positive association between this chemokine and CD206 expression in B78chOVA TAMs (**Fig. 3I**), resulting in a significant difference (again ∼50%) in CXCL9 expression in CD206+ vs. CD206- TAMs (**Fig. 3J**) in the WT mice. This finding was substantiated in another subcutaneously injected tumor model MC38chOVA and the spontaneous breast tumor model PyMTchOVA, both with lower overall CXCL9 positivity in the absence of OT-I adoptive transfer, but a consistent ∼50% or more difference between the CD206+ and CD206- groups (**Supplementary Fig. S3E**). CD206+ monocytes also showed higher CXCL9 expression, compared to CD206- counterparts (**Supplementary Fig. S3F**), but CXCL9+ monocytes were only 1/4^th^ as abundant as CXCL9+ TAMs in the B78chOVA TME, thus limiting their relative role in myeloid CXCL9 production (**Supplementary Fig. S3G**). We therefore sorted CD206+ vs. CD206-TAMs from B78chOVA tumors (d14 post tumor injection without OT-I treatment) and interrogated their relative effects on in vitro activated CD8 T cell transmigration in a 3h window. Consistent with their CXCL9 expression, the CD206+ but not the CD206- TAMs induced enhanced transmigration over no TAM-added controls. (**Fig. 3K**). Since CD206 TAM-depleted tumors still had small numbers of lymphocytes, we compared their levels of CXCR3 at the transcript level, which reflects receptor-ligand engagement avoiding the confounding effect of receptor internalization(*21*), and found that *Cxcr3* expression was markedly lower in the DTx treated condition in all the lymphocyte subsets (**Fig. 3L, Supplementary Fig. S3H**). Taken together, these data point to the role of CD206+ TAMs in the recruitment of CXCR3-expressing lymphocytes to the TME.

Lymphocytes are well-established as key producers of cDC1-formative chemokines FLT3L (*18*) and XCL1(*22*). Therefore, given the loss of cDC1s in concert with lymphocytes with this depletion regime, we probed for these chemokines in the intratumoral lymphocytes in control vs. CD206- depleted dataset. This demonstrated that both *Flt3l* and *Xcl1* (**Fig. 3M, Supplementary Fig. S3I**) transcripts were markedly reduced in the NK cells and CD8 T cells in the DTx-treated condition. These changes on a per cell basis, in addition to the overall decrease in CD8 T cells and NK cells are consistent with the loss of cDC1s in the TME, as a result of the disruption of the CD8:NK:cDC1 module (*18*).

### Depletion of CD206+ TAMs thwarts CD8 T cell mediated anti-tumor immunity in mice

Given our finding that CD206+ ‘M2-like’ macrophages are critical to the organization a key node of anti-tumor immunity, we asked whether they were necessary for successful CD8 T cell mediated tumor regression. To test this, we used a MC38chOVA model where an adoptive transfer of OT-I T cells that results in efficient tumor control(*17*) (**Supplementary Fig. S3J**). We confirmed first that reporter expression in these tumors followed largely the same pattern as the B78chOVA tumors, with substantial expression only in TAMs and cDC2s, albeit at lower levels due to the overall lower CD206+ fraction, and little to no expression in neutrophils and monocytes in this model. (**Supplementary Fig. S3K, L**). Importantly, lymphocytes and cDC1s again showed no reporter expression (**Supplementary Fig. S3K, L**). As with the B78chOVA model, we applied early and late depletion regimens to the MC38chOVA tumors, first without the addition of OT-I T cells, to assess differential effects on early establishment and maintenance of immune cells without the confounding variable of tumor regression (**Fig. 4A**). Some differences were observed compared to the B78chOVA model, including an overall maintenance of TAM abundance in late and only a modest (∼25%) decline in early depletion (**Supplementary Fig. S3N, R**) despite robust depletion of the CD206+ populations (**Fig. 4B, E**). Indeed, following reporter expression patterns, the CD206+ subsets were depleted robustly in TAMs and cDC2s and modestly in monocytes and neutrophils in both the late (**Fig. 4I**) and early (**Fig. 4J**) depletion regimens. We also noted other variations namely increased monocyte abundance in early but not late depletion (**Supplementary Fig. S3O, S**) and neutrophil enrichment in both regimens (albeit with much lower effect size in late depletion; **Supplementary Fig. S3Q, U**). As in the case of B78chOVA tumors, CD206+ cDC2s were depleted (**Fig. 4I, J**) but no change was detected in overall cDC2 abundance (**Supplementary Fig. S3P, T**) with both early and late DTx treatment. Importantly, there was once again a robust indirect loss of cDC1s and CD8 T cells specifically under the early but not the late depletion regimen (**Fig. 4C, D, F, G**). When evaluated further by directly measuring total number of cells per g of tumor, we observed once again the direct depletion of CD206+ TAMs, indirect increase in neutrophils and the indirect loss of cDC1s and CD8 T cells, but not NK cells (**Supplementary Fig. S3U**) in MC38chOVA tumors with early DTx treatment. With confirmation of this key indirect effect of CD206+ TAM depletion in the MC38chOVA model, we treated subcutaneous MC38chOVA tumors in WT and DTR mice with OT-Is and concomitant early DTx administration. With the prediction that depletion of CD206+ TAMs would thwart the tumor control ability of OT-Is, we tracked changes in tumor size and indeed observed significantly reduced OT-I-mediated tumor control of MC38chOVA tumors (**Fig. 4H**) in the DTR group.

**Fig. 4:**
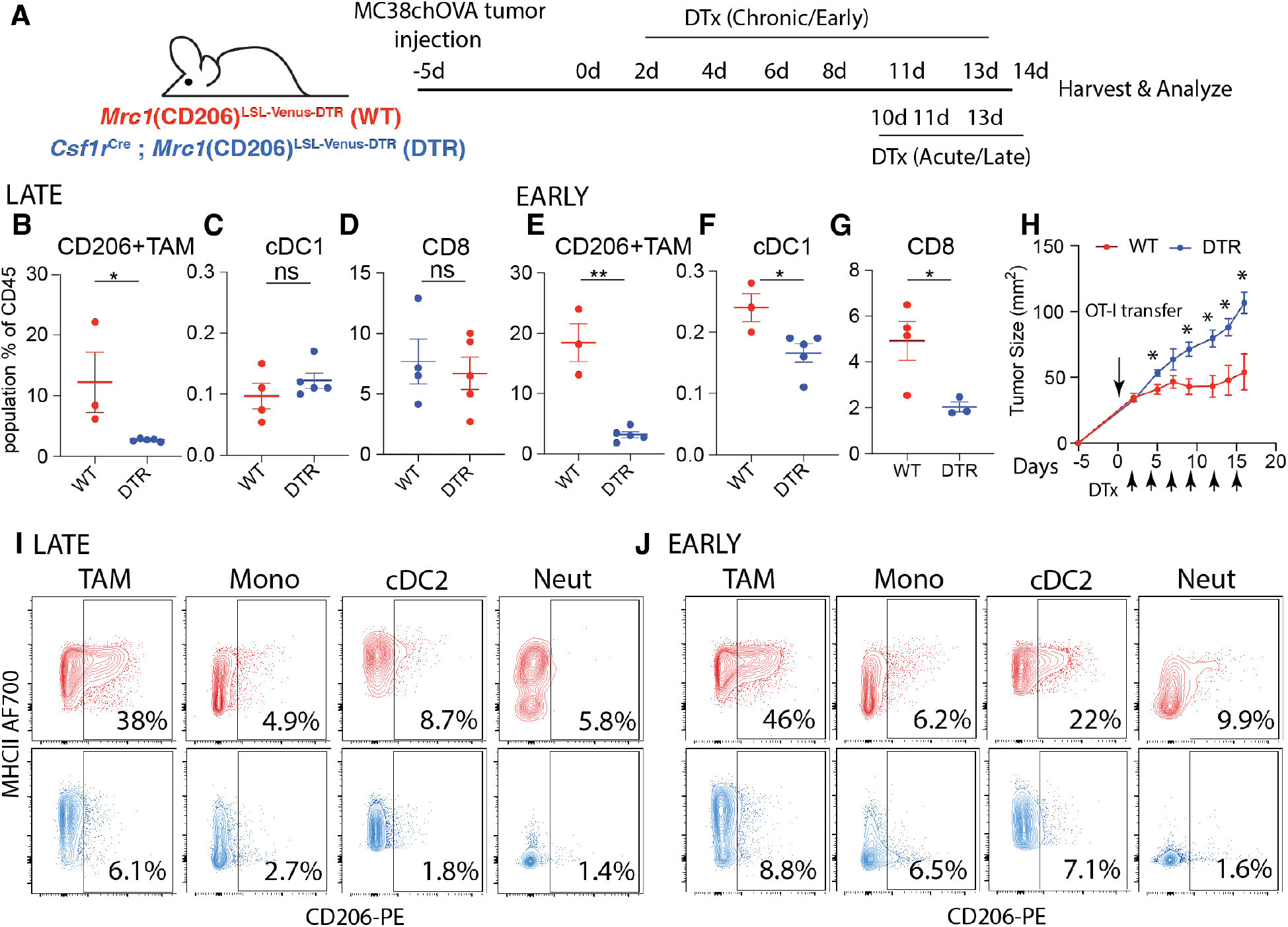
CD206+ TAM depletion disrupts the cDC1:CD8 module and attenuates T cell-mediated tumor control in an immune-responsive tumor model: **(A)** Schematic representation of the experimental setup for early and late CD206+ TAM depletion in MC38chOVA tumors using *Csf1r*^Cre^; *Mrc1*^LSL-Venus-DTR^ mice; Relative abundance of **(B, E)** CD206+ TAMs, **(C, F)** cDC1s and **(D, G)** CD8 T cells as a percentage of CD45+ cells with late and early depletion regimens respectively; **(H)** tumor growth kinetics of MC38chOVA tumors in WT and DTR mice with DTx treatment beginning 2d post OT-I adoptive transfer at Day 0; Representative flow cytometry plots showing CD206 vs. MHCII expression in different myeloid subsets in WT (red) and DTR (blue) mice in the (**I**) late and (**J**) early depletion regimens; **p<0.01, *p <0.05, ns = no significance by Mann-Whitney U test t-tests.

### CD206+ Mono/Mac signatures associate with anti-tumor immunity in human cancers

The data presented thus far provided substantial evidence of a context in which CD206+ populations of Mono/Macs were in fact positive contributors to reactive anti-tumor immunity in mice, rather than being simply immunosuppressive. Consistent with this understanding, we found that higher levels of *MRC1* RNA alone correlated with slightly better survival from patient data rather than worse in a large cohort curated from The Cancer Genome Atlas (TCGA) (*23*) (**Fig. 5A**). We also sought to determine whether the revealed relationships between CD206+ TAMs, CXCL9 and the cDC1:CD8:NK module in our study might similarly extended to human disease. To do so, we first applied differential gene expression (DGE) analysis of the Ctrl vs. DTx treated Mono/Mac populations (**Fig. 5B, C**) (excluding neutrophils, cDC1, cDC2 and lymphocyte subsets) from our scRNASeq dataset (**Fig. 3B**). We then used the top 10 DEGs (by average log fold change and having an adjusted p-value <0.01) to create CD206 ‘Replete’ and ‘Depleted’ gene signatures (**Fig. 5D**). The former are DEGs associated with the presence of CD206+ populations and not only included *C1qa, Cxcl9, Apoe,* but also several MHC-II related genes (**Fig. 5D, E**), consistent with flow cytometry data on C1q TAM, CD206, CXCL9 and MHC-II expression described above. The Depleted signature contains genes differentially expressed in macrophages that remain post CD206+ Mono/Mac depletion and included *Il1b*, *S100a8*, along with *Spp1* (**Fig. 5D, E**). Even though we obtained these gene signatures based upon depletion of Mono/Macs using the prominent “M2” marker CD206, both M1 and M2-associated genes were differentially upregulated in Replete signature (**Fig. 5D**), reiterating the lack of coordination among such markers when studied in vivo.

**Fig. 5:**
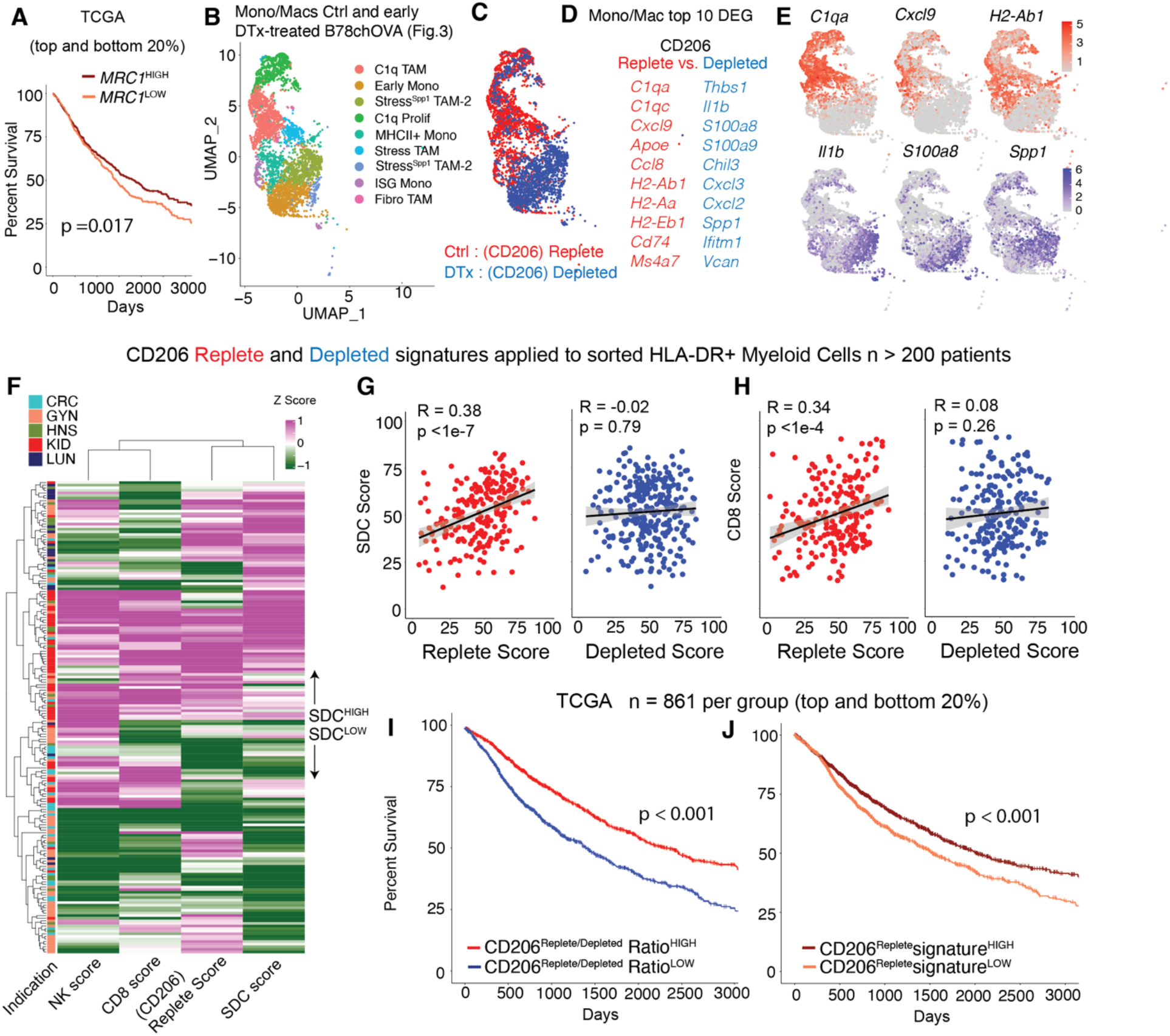
CD206^Replete^ Mono/Mac signature associates with anti-tumor immunity in human cancers: **(A)** Kaplan-Meier survival curves of patients in TCGA grouped by the expression of *MRC1* gene; **(B)** UMAP representation of the Mono/Mac subsets and **(C)** overlay of the CD206 Replete (Ctrl) and Depleted (DTx) groups on the UMAP from spatial scSeq described in Fig. 3; **(D)** Top 10 genes from Differential Gene Expression (DGE) of Mono/Macs in the Ctrl vs. DTx treated conditions, which were used to generate CD206 Replete and CD206 Depleted Mono/Mac signature scores **(F)** Heatmap of z-scored CD206^Replete^, CD8, NK and Stimulatory dendritic cell (SDC) score, calculated from sorted immune compartments, as previously described (*23*); Scatter plots of the Myeloid-specific CD206 Replete and Depleted score per patient with the **(G)** stimulatory dendritic cell (SDC) score and (**H**) CD8 T cell score (Pearson R and p value for the null hypothesis that there is not a correlation are noted); Kaplan-Meier survival curves of patients grouped by the value of the (**I**) CD206^Replete^: CD206^Depleted^ signature ratio and (**J**) CD206^Replete^ signature in TCGA, p values for the log-rank test are noted in (A, I, J).

Using these signatures, we queried a previously described immune compartment-specific bulk RNA-Seq data derived from sorted HLA-DR+ myeloid (to capture Mono/Macs and DCs), T and total live cells from >200 human tumor biopsies(*23*) (**Fig. 5F, G**) belonging to five common solid tumor indications (CRC: Colorectal Cancer, GYN: Gynecological Cancer, HNSC: Head and Neck Squamous Cell Carcinoma, KID: Kidney Cancer; LUNG: Lung Carcinoma). Given our finding that the CD206+ CXCL9-expressing TAMs recruit CXCR3-expressing cDC1-supportive lymphocytes, we predicted that the CD206 “Replete” but not the “Depleted” signature in the myeloid compartment would associate with previously established CD8, NK and stimulatory cDC1s (stimulatory dendritic cell or SDC) gene signatures (*10*) (*23*). Indeed, the Replete signature, but not the Depleted signature correlated significantly with those of each component of the tumor-reactive immune module (**Fig. 5F**) of SDCs (**Fig. 5G**), CD8 T cells (**Fig. 5H**) and NK cells (**Supplementary Fig. 3W**). Given that it is by now well-established that SDCs are associated with survival (*10*), we also queried whether the relative abundance of CD206 Replete Mono/Macs (i.e., CD206^Replete^/CD206^Depleted^ ratio) was correlated with better survival in patients. In the TCGA dataset, we observed a large (∼20% in 5-year survival) and significant shift in survival for patients biased towards the CD206^Replete^ Mono/Mac signature (**Fig. 5I**). Indeed, patients scoring high on the CD206^Replete^ Mono/Mac signature alone were also found to have significantly better survival but with a reduced effect size (**Fig. 5J**) as compared to the ratio. Among specific indications, the Replete/Depleted signature ratio was associated with overall survival in Lung, Liver, Pancreatic, Bladder, Kidney Cancer and Melanoma (**Supplementary Fig. S3X**). Thus, contrary to the simplistic labeling of CD206 expressing macrophages as immunosuppressive, this data establishes contexts in which these Mono/Macs are a critical organizing fulcrum for the reactive archetype of NK cells, cDC1s and CD8 T cells. Taken together, these data contribute to a nuanced understanding of the context-dependent role of TAMs in the TME, necessary to rationally design next-generation myeloid-targeting immunotherapies in cancer.

## Discussion

cDC1s have been previously linked to FLT3L and XCL1 producing NK cells and activated CD8 T cells(*18, 22*), and this network represents one module of immunity that predisposes to immune checkpoint blockade response (reviewed in(*24*)). The same CD8 T cells in turn may be recruited and expanded by chemokines and antigen-presentation by cDC1s, creating a virtuous feedback loop for anti-tumor immunity. It was however, previously unexplored how specific macrophage subsets support or thwart this anti-tumor archetype. Here, we demonstrate that CD206 expression in macrophages is robustly correlated with their expression of CXCL9 and these macrophages play a critical role in initiating the assembly of the cDC1:NK-CD8 anti-tumor reactive immune archetype in tumors.

This work is the latest in a series of publications (*3, 5, 12*). that force re-evaluation of the prevalent but insufficient M1/M2 classification of macrophages in tumors. Notably, CD206 expression is still often used to categorize macrophages as immunosuppressive and ‘M2-like’, even though strong in vivo data supporting this assertion is lacking. Here, we show that CD206 should not be used as an unqualified indicator of immunosuppressive function. Indeed in the context of ongoing anti-tumor responses studied here in the early depletion setting, these TAMs are crucial for effective recruitment of critical immune cells.

When thinking about these revealed functions of TAMs expressing CD206, we also note that this study specifically found them critical in early T cell recruitment. However, our work and others have shown that some mature TAMs which include those that express CD206 may also be involved in coupling with CD8 T cells and promote T cell exhaustion (*11, 25*). Thus, TAMs may have distinct phenotypes and functions depending on the immunological state of the tumor – perhaps reflected in the early and late depletion conditions shown here. Future studies to understand this balance of pro and anti-tumor effects of TAMs is critical. At present, one should not simplistically take our study to indicate that CD206+ macrophages are universally favorable for anti-tumor immunity. However, the M1/M2 dichotomy—and particularly a version that equates CD206 with pro-tumoral functions—appears to be a misleading lens through which to view macrophage functional heterogeneity.

Other recent data using *CXCL9* versus *SPP1* gene expression to functionally classify macrophages in human tumors (*12*) as anti- or pro-tumor respectively, are aligned with our findings. In our earlier studies of *SPP1* in macrophages (*3*), these were observed in human tumors, likely embedded within an Arg1 (Stress) TAM subset in mice, and here they only emerged as a distinct cluster due to their disproportionate enrichment post depletion. As we have previously noted(*3*), these non-CD206 expressing ‘Stress’ macrophages are distinctly glycolytic, express *Hif1α*, and are likely the cells that have previously been defined as hypoxic macrophages (*26, 27*) and now associated with poor patient outcomes.

Studies prior to ours and using more universal Mono/Mac depletion have also variably reported compensatory neutrophil influx when depleting cells of monocytic origin in tumors (*28–30*). Our data also shows an increase in neutrophils in the early CD206-gated depletion condition, but a lack of similar influx in the late depletion regimen. One interpretation of our data is that a microenvironment-dependent opportunistic filling of the early myeloid niche by neutrophils takes place in the absence of sufficient Mono/Macs and the reactive immune components. While further studies may uncover key nodes of this balance of myeloid populations, our results show that the compensatory neutrophils do not contribute to the reduction of CXCR3-dependent lymphocyte recruitment (*20*), which is the primary driver leading to the loss of the key tumor-reactive archetype.

One key success of our study is the ability to differentially target subsets of TAMs within the TME. Prior to our work, many questions regarding the specific role of TAMs have remained obscured or unanswered partly owing to the lack of sufficiently specific and penetrant tools to manipulate them in vivo. Commonly used methods, while useful lack sufficient specificity, including the depletion of all monocytes and monocyte-derived dendritic cells (CSF1R blocking antibody;(*31–33*)) and the depletion of all phagocytic cells and arrest of neutrophils (Clodronate; (*34*)). In this context, the novel conditional CD206 reporter introduced here—paired with *Csf1r*-Cre to avoid depleting other non-myeloid cells that express CD206—provides a more selective marking and depletion tool for CD206+ TAMs, with a further potential to target various subpopulations by altering the Cre driver alleles.

Overall, our results indicate that even this subset-dependent depletion of Mono/Mac populations may not be prudent in all contexts. To this extent, while anti-CSF1R antibodies have failed to show benefits in clinical trials (*35*), other strategies using for example, drugs that modulate specific subsets such as those expressing TREM2 or trigger others by engaging TREM1 may prove more surgical(*36*). Systematically dissecting the role of individual TAM subtypes will continue to be crucial to deciphering their context-dependent and complex roles in the TME, with a view towards harnessing them for better immunotherapy outcomes.

## Funding

National Institutes of Health Grants: NIH R01CA197363 and NIH R37AI052116

AR was supported by a Cancer Research Institute Postdoctoral Fellowship (CRI2940)

KHH was supported by an American Cancer Society and Jean Perkins Foundation Postdoctoral Fellowship

NFK was supported by the Cancer Research Institute / Merck Postdoctoral Fellowship (CRI4546)

We thank members of the Krummel lab for their inputs to the manuscript.

## Author Contributions

Conceptualization: AR, MFK

Experimentation: AR, KK, IZL, NFK

ZipSeq: KHH, AR

Human tumor data curation and analysis: TC, AJC, BS

Writing: AR, MFK

Supervision: MFK

## Declaration of Interests

The authors declare no competing interests

## Data and materials availability

Relevant data will be made publicly available before publication in its final form. Meanwhile, data will be available upon reasonable request, please contact the authors directly.

## List of Supplementary Materials

Materials and Methods

Fig. S1-S3

## Supplementary Materials for

### Materials and Methods

#### Mice

All mice were treated in accordance with the regulatory standards of the National Institutes of Health and American Association of Laboratory Animal Care and were approved by the UCSF Institution of Animal Care and Use Committee. *Mrc1*(CD206)^LSL-Venus-DTR^ mice in the C57BL6/J background were custom-generated from Biocytogen Inc. and then maintained heterozygous (bred to C57BL6/J wild type mice) at the UCSF Animal Barrier facility under specific pathogen-free conditions. C57BL6/J, C57BL6/J CD45.1 (B6.SJL-Ptprc^a^ Pepc^b^/BoyJ), OT-I (C57BL/6-Tg(TcraTcrb)1100Mjb/J), Csf1r^Cre^ (C57BL/6-Tg(Csf1r-cre)1Mnz/J) mice were purchased for use from Jackson Laboratories and maintained in the same facility in the C57BL6/J background. For adoptive transfer experiments, CD45.1^het^; OT-I^het^ (denoted simply as CD45.1; OT1) mice were used. Mice of either sex ranging in age from 6 to 14 weeks were used for experimentation.

#### Depletion of select immune cell populations

For depletion of CD206-expressing macrophages, 500ng (20ng/g body weight, assuming an average 25g weight for each mouse) diphtheria toxin (DTx; List Biological Laboratories) in 100µL 1X PBS was injected intraperitoneally into each mouse – for both *Csf1r*^Cre^; *Mrc1*(CD206)^LSL-Venus-DTR^ (DTR) and *Mrc1*(CD206)^LSL-Venus-DTR^ (WT) groups - at every time point. For the early depletion regime, injections were started 2 days after adoptive transfer of T cells and continued every 2-3 days till endpoint, while for the late depletion regime, injections began at d10 after T cell injection and continued till endpoint. For testing the effects of DTx in tumor-free tissue, similar dosing of DTx as the early depletion regime was implemented without tumor injection, and the skin (ectopic tumor site) and skin-draining lymph nodes were isolated for analysis. Mice were found to be healthy and without frank health issues with 6 doses of 500ng DTx (early depletion regime), but were monitored nevertheless throughout the experiment, as per IACUC guidelines.

For depletion of neutrophils, mice were treated with 200µg/dose of anti-Ly6G antibody (Clone 1A8, InvivoMAb) in PBS intraperitoneally every 2-3 days starting one dose after the beginning of DTx treatment and coincident with DTx treatment thereafter. Control mice were similarly treated with the corresponding isotype control antibody (Clone 2A3, InvivoMAb).

#### Mouse tumor digestion and flow cytometry

Tumors from mice were processed to generate single cell suspensions as described previously(*18*). Briefly, tumors were isolated and mechanically minced on ice using razor blades, followed by enzymatic digestion with 200 μg/mL DNAse (Sigma-Aldrich), 100U/mL Collagenase I (Worthington Biochemical) and 500U/mL Collagenase IV (Worthington Biochemical) for 30 min at 37°C while shaking. Digestion was quenched by adding excess 1X PBS, filtered through a 100μm mesh, spun down and red blood cells were removed by incubating with RBC lysis buffer (155 mM NH_4_Cl, 12 mM NaHCO_3_, 0.1 mM EDTA) at room temperature for 10 mins. The lysis was quenched with excess 1X PBS, spun down and resuspended in FACS buffer (2mM EDTA + 1% FCS in 1X PBS) to obtain single cell suspensions. Similarly, tumor draining lymph nodes (dLN) were isolated and mashed over 100μm filters in PBS to generate single cell suspensions. For counting absolute numbers of cells, CountBright Absolute Counting Beads were added to the cell suspensions prior to staining, while noting the total weight of the tumor and the fraction of the total tumor cell digest used for staining.

For each sample, 2.5-3 million cells/sample were stained in a total of 50µL of antibody mixture for flow cytometry. Cells were washed with PBS prior to staining with Zombie NIR Fixable live/dead dye (1:500) (Biolegend) for 20 min at 4°C. Cells were washed in FACS buffer followed by surface staining for 30 min at 4°C with directly conjugated antibodies diluted in FACS buffer containing 1:100 anti-CD16/32 (Fc block; BioXCell) to block non-specific binding. Antibody dilutions ranged from 1:100-1:400, optimized separately. After surface staining, cells were washed again with FACS buffer. For intracellular staining, cells were fixed for 20 min at 4°C using the IC Fixation Buffer (BD Biosciences) and washed in permeabilization buffer from the FoxP3 Fix/Perm Kit (BD Biosciences). Antibodies against intracellular targets were diluted in permeabilization buffer containing 1:100 Fc Block and cells were incubated for 30 min at 4°C followed by another wash prior to readout on a BD LSRII or Fortessa Cytometer.

#### Processing and flow cytometry analysis of other mouse organs

To phenotype cells from lymphoid organs, inguinal, axillary and brachial (tumor-draining) lymph nodes were isolated, pried open with tweezers (lymph nodes) or cut into small pieces (spleen) and digested with the same digestion cocktail as above, intermittently pipetting with cut P1000 pipette tips to enhance mechanical digestion. The resulting suspensions were then filtered using 100µm filter, washed with 1X PBS to generate single cell suspensions. For splenic digests, RBC lysis was performed as described above before staining for flow cytometry.

For lung digests both lobes were isolated, cut into small pieces with scissors and minced by using gentleMACS dissociator (Miltenyi Biotec) in RPMI. Next, the mixture was spun down and resuspended in the digestion mixture described above and allowed to digest with shaking at 37°C for 20 mins, following which, the remaining tissue was either minced again using the gentleMACS dissociator and/or directly mashed over a 100µm filter in FACS buffer to generate a single cell suspension, ready to be processed for staining and flow cytometry.

Skin digestion was done as previously described(*37*). Briefly, mice were shaved and depilated prior to removal of dorsal skin. The skin was then rid of fat, minced with scissors and razor blade in the presence of 1 ml of digest media (2 mg/ml collagenase IV (Roche), 1 mg/ml hyaluronidase (Worthington), 0.1 mg/ml DNase I (Roche) in RPMI-1640 (GIBCO). The minced skin was then moved to a 50 ml conical with 5 ml additional digest solution and incubated at 37°C for 45 min with shaking and intermittent vortexing before being washed and passed through a 70μm strainer prior to staining.

#### Flow cytometry Data Analysis

Analysis of flow cytometry data was done on FlowJo and later plotted on GraphPad Prism or R. Relative MFI of the Venus reporter was calculated by subtracting the background average MFI of the same channel in WT samples from those in each DTR sample. For analysis of a shift in relative abundance of a population x (Fig. 2), the log_2_ (% x of CD45 in WT/ % x of CD45 in DTR) was calculated and plotted as a heatmap, such that positive values indicate depletion and negative values indicate enrichment.

#### Tumor injections and adoptive transfer of CD8 T cells into tumors

The B78chOVA and MC38chOVA cancer cell lines, as previously described(*11, 18*), were generated by incorporating the same mcherry-OVA construct used to establish the PyMTchOVA spontaneous mouse line(*38*). For tumor injections, the corresponding cells were grown to near confluency (cultured in DMEM with 10% FCS (Benchmark) and 1% PSG (Gibco)) and harvested using 0.05% Trypsin-EDTA (Gibco) and washed 3x with PBS (Gibco). The number of cells to be injected per mouse was resuspended in PBS to a final volume of 50μL per injection. The suspension was injected subcutaneously into the flanks of anesthetized and shaved mice. Tumors were allowed to grow for 14–21 days unless otherwise noted, before tumors and tumor-draining lymph nodes were harvested for analysis. CD8 T cells were isolated from CD45.1;OT-1;Cd69-TFP mice using the EasySep Negative Selection Kit (Stem Cell Bio), resuspended in 1X PBS at 10X concentration 100µL was injected into each tumor-bearing mice. For B78chOVA 1 million and for MC38chOVA tumors, 200,000 CD8 T cells were injected retro-orbitally into each mouse either 5d (B78chOVA), 7d (MC38chOVA) post tumor injection. Tumor measurements were done by measuring the longest dimension (length) and approximately perpendicular dimension (width) using digital calipers, rounded to one decimal place each. For experiments using the transgenic PyMTchOVA strain, mammary tumor-bearing females in the age range of 15 to 24 weeks were used when mice developed at least 2 palpable tumors.

#### Spatial single cell RNA Sequencing and Analysis

Spatial scSeq of immune cell populations at the tumor edge, interface and interior zones was performed using ZipSeq, as previously described(*11*), with the additional condition of DTx treatment integrated into the dataset. Briefly, B78chOVA tumors subcutaneously grown in Csf1rCre; CD206^LSL-Venus-DTR^ mice d12 post adoptive transfer of 1 million CD2dsRed; OT-I CD8 T cells with (DTx) and without (Control) DTx treatment (early depletion regime) were harvested and sliced into 160µm slices using a Compressotome (Precisionary Instruments VFZ-310-0Z). Imaging, spatial barcoding, subsequent digestion, sorting, encapsulation (10X Genomics) and library construction, CellRanger processing and alignment were performed as described previously(*11, 19*). The two separate sequencing runs (Control and DTx) were assembled and integrated into a single data structure using Harmony(*39*). The final object underwent scaling and then scoring for cell cycle signatures (S and G2M scores as computed using Seurat’s built-in CellCycleScoring function. The object then underwent regression for cell cycle effects (S and G2M score as described in the Seurat vignette) and percent mitochondrial reads before PCA.

Relative abundance from scSeq data was calculated by: log2 (% of each cluster (cell type) within a tumor region (Edge, Mid, Inner) in the Ctrl / (% of the same cluster in the same region in the DTx treated group), thereby yielding positive values for depletion and negative values for enrichment. While abundances were calculated with the broad clusters from the overall object, the lymphoid clusters were isolated to a separate object, re-clustered to further probe for individual gene expression (*Cxcr3, Flt3l, Xcl1*) in the resulting subsets.

#### Transwell Assay of CD8 T cell migration

For transwell assays, subcutaneously injected B78chOVA tumors grown for 14 days and then harvested, digested, and sorted for CD206+ vs. CD206- TAMs. 3 days before the sort, CD8 T cells from a B6 mouse were harvested and stimulated in vitro with anti-CD3/anti-CD28 Dynabeads (Thermo Fisher) for 24h, taken off the beads and rested in 10U/mL IL-2 for an additional 48h to produce effector-like CD8 T cells. Post-sort, 500,000 activated T cells were plated in 75µL T cell media (RPMI + 10% FCS + 50µM β - marcaptoethanol) on top of a 5µm transwell insert (Corning), allowed to settle for 30mins and subsequently, 10,000 sorted CD206-, CD206+ TAMs or no TAMs were added to the bottom well to induce T cell migration. Cells at the bottom were collected at 3h, mixed with CountBright absolute counting beads, stained and analyzed by flow cytometry to quantify the number of CD8 T cells migrated. Total number of CD8 T cells migrated in each condition was normalized to the average number of cells migrated in the no TAM condition.

#### Human tumor samples

All tumor samples were collected with patient consent after surgical resection under a UCSF IRB approved protocol (UCSF IRB# 20-31740), as described previously(*23*). In brief, freshly resected samples transported in ice-cold DPBS or Leibovitz’s L- 15 medium before digestion and processing to generate a single-cell suspension. The five most well-represented cancer indications in this collection were included in the cohort: Colorectal cancer (CRC), gynecological cancers (GYN), head and neck cancer (HNSC), kidney cancer (KID), lung cancer (LUNG). Clinical data including survival of patients were obtained through regular clinical follow-up at UCSF.

#### Transcriptomic analysis of human tumors

All tumor samples were collected under the UCSF Immunoprofiler project as described(*23*). Briefly, tumor samples were thoroughly minced with surgical scissors and transferred to GentleMACS Tubes containing 800 U/ml Collagenase IV and 0.1 mg/ml DNase I in L-15/2% FCS per 0.3 g tissue. GentleMACS Tubes were then installed onto the GentleMACs Octo Dissociator (Miltenyi Biotec) and incubated for 20 min (lymph node) or 35 min (tumor) according to the manufacturer’s instructions. Samples were then quenched with 15 mL of sort buffer (PBS/2% FCS/2mM EDTA), filtered through 100μm filters and spun down. Red blood cell lysis was performed with 175 mM ammonium chloride, if needed. Freshly digested tumor samples were sorted by FACS into conventional T cell, Treg, Myeloid, tumor and in some cases, stromal compartments and bulk RNA-seq was performed on sorted cell fractions. mRNA was isolated from sorted fractions and libraries were prepared using Illumina Nextera XT DNA Library Prep kit. The libraries were sequenced using 100bp paired end sequencing on HiSeq4000. The sequencing reads we aligned to the Ensembl GRCh38.85 transcriptome build using STAR(*40*) and gene expression was computed using RSEM(*41*). Sequencing quality was evaluated by in-house the EHK score, where each sample was assigned a score of 0 through 10 based on the number of EHK genes that were expressed above a precalculated minimum threshold. The threshold was learned from our data by examining the expression distributions of EHK genes and validated using the corresponding distributions in TCGA. A score of 10 represented the highest quality data where 10 out of 10 EHK genes are expressed above the minimum threshold. The samples used for survival analysis and other gene expression analyses had an EHK score of greater than 7 to ensure data quality. Ensemble gene signatures scores were calculated by converting the expression of each gene in the signature to a percentile rank among all genes and then determining the mean rank of all the genes in the signature (*17*). The corresponding gene list for obtaining the stimulatory dendritic cell score is as described before(*10*).

#### TCGA analyses

Survival analyses using the TCGA dataset was performed using the TCGA sub-cohort described in(*23*). Briefly, tumor RNAseq counts and TPM along with curated clinical data for 13 cancer types (BLCA, COAD, GBM, GYN (grouping OV, UCEC and UCS), HNSC, KIRC, LIHC, LUAD, PAAD, SARC and SKCM) was filtered down to include primary solid tumors and metastatic samples only, to parallel the IPI cohort samples. This reduced the TCGA sample set to 4341 tumor samples. CD206^Replete^ gene scores were generated by first normalizing (using percentiles) the expression values of each gene composing the signature across all patients, followed by averaging these normalized values for each patient. The same method was used for deriving CD206^Depleted^ gene scores and we then calculated the ratio of CD206^Replete^/CD206^Depleted^ gene scores by dividing each score value for each patient. For survival analysis, patients were split into either CD206^Replete^ gene score ^HIGH^ vs ^LOW^ (top/bottom 20% respectively, n=861) or (CD206^Replete^:CD206^Depleted^ gene signature ratio)^HIGH^ vs (CD206^Replete^:CD206^Depleted^ gene signature ratio)^LOW^ (top/bottom 20% respectively, n=861) and analyzed using a log-rank test.

#### Two-photon imaging of tumor slices

Tumor slices (adjacent to the ones used for spatial barcoding by ZipSeq) were fixed in 2% paraformaldehyde (PFA; Sigma), washed and left overnight in 1X PBS before imaging on a custom-made 2-photon microscope as previously described(*10*) to visualize the Venus reporter and CD2dsRed marked CD8 T cells and fibrous collagen by second harmonic generation (SHG). Dual laser excitations at 800nm and 950nm were used to excite the requisite fluorophores.

#### Statistical Analysis

Statistical analysis was done in GraphPad Prism or in R. For testing null hypothesis between two groups, either Student’s t tests and or the non-parametric Mann-Whitney U tests were used, depending on the number and distribution of data points. Likewise, for testing null hypotheses among 3 or more groups, ANOVA or non-parametric tests were performed, followed by post-hoc test, correcting for false discovery rates (threshold = 0.05) in multiple comparisons. Unless otherwise mentioned, data are representative of at least 2 independent experiments.

## Supplementary Figures and Figure Legends

**Fig. S1:**
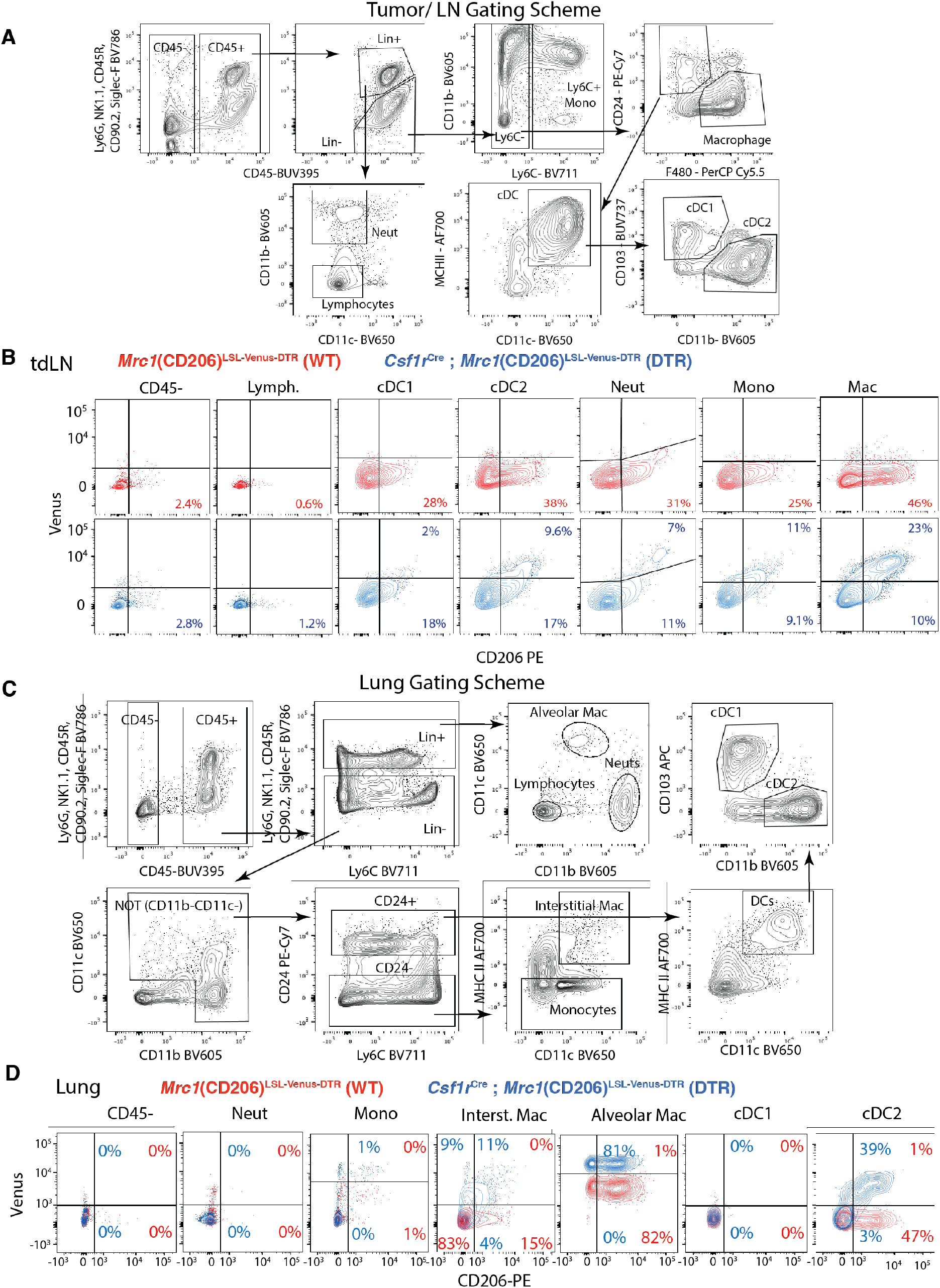
Representative flow cytometry gating scheme to identify myeloid cells and lymphocytes from **(A)** tumor and tdLN and **(C)** lung;Flow cytometry plots showing reporter (Venus) and CD206 expression in different immune cells in (**B**) d18 B78chOVA tdLN and (**C**) lung in WT (red; *Mrc1*^LSL-Venus-DTR^) and DTR (blue; *Csf1r*^Cre^; *Mrc1*^LSL-Venus-DTR^) mice.

**Fig. S2:**
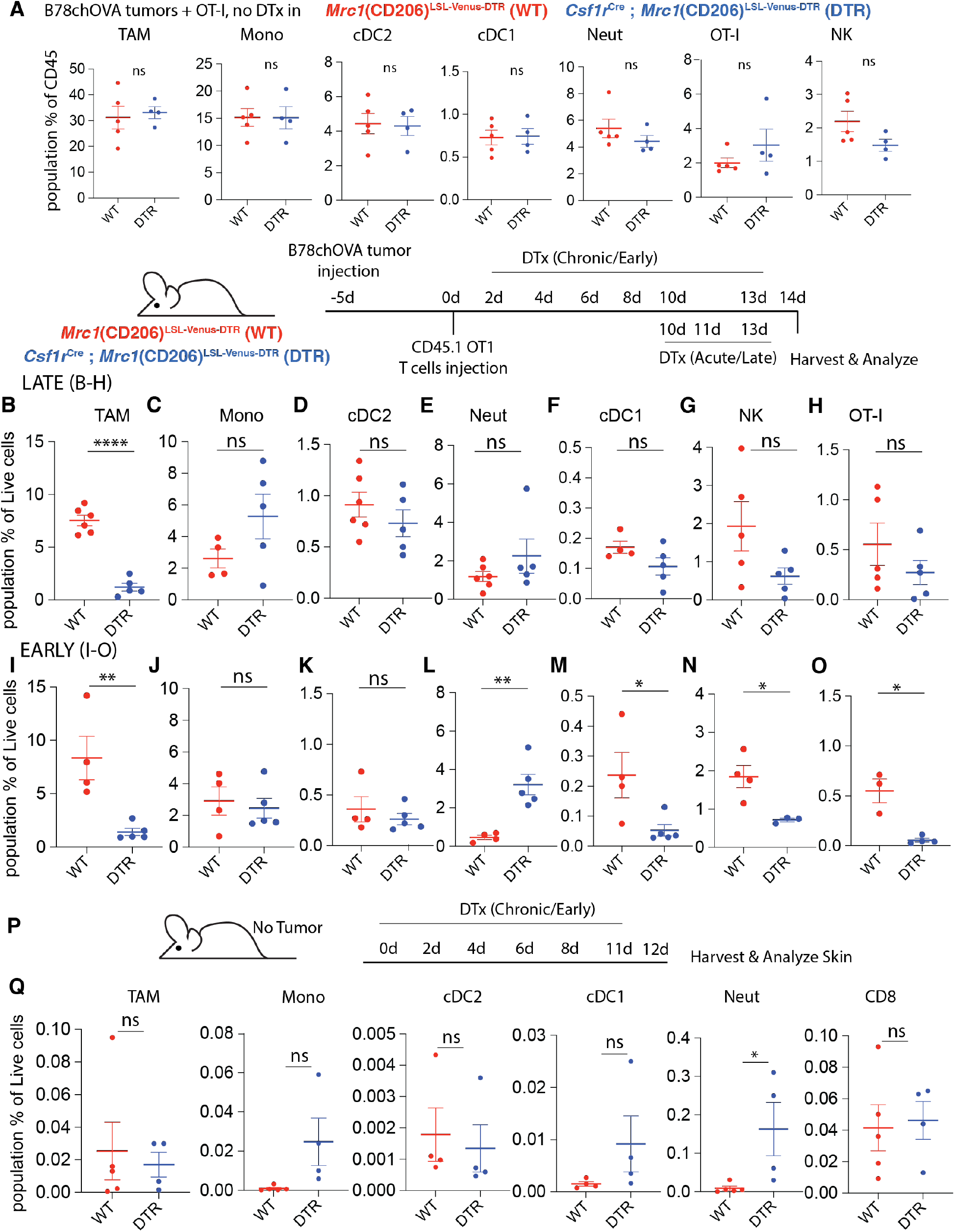
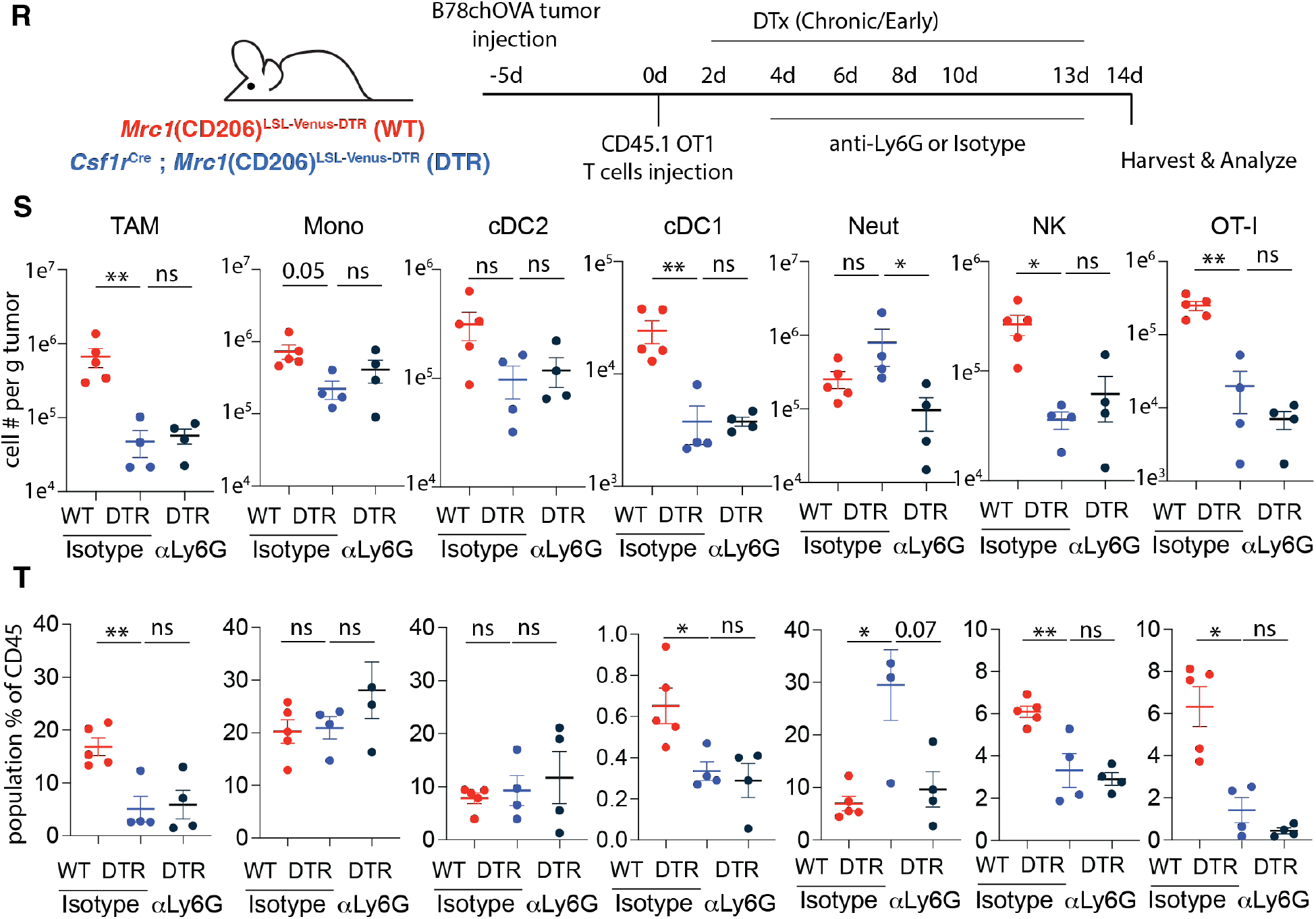
**(A)** Relative abundance of different immune populations as a percentage of CD45+ cells in *Mrc1*^LSL-Venus-DTR^ (WT) and *Csf1r*^Cre^; CD206^LSL-Venus-DTR^ (DTR) mice with B78chOVA tumors (day-5), OT-I adoptive transfer (day 0) and harvest at d14 without DTx administration; Schematic representation of the experimental setup for tumor injection, OT-I T cell adoptive transfer, early and late diphtheria toxin administration and analysis; Relative abundance of different immune populations as a percentage of live cells with **(B-H)** late and **(I-O)** early depletion regimens. (**P**) Schematic representation of the experimental setup for analysis of skin in *Mrc1*^LSL-Venus-DTR^ (WT; red) and *Csf1r*^Cre^; *Mrc1*^LSL-Venus-DTR^ (DTR; blue) mice with DTx administration; **(Q)** Relative abundance of different immune populations in the skin as a percentage of live cells; (**R**) Schematic representation of the experimental setup for B78chOVA tumor injection, OT-I T cell adoptive transfer, and early diphtheria toxin administration with either isotype control or anti-Ly6G antibody treatment and analysis; Abundance of different immune populations as (**S**) cells per g of tumor and **(T)** percentage of CD45+ cells in WT and DTR mice. ****p<0.0001, **p<0.01, *p <0.05, ns = no significance by Student’s t-tests or Mann-Whitney test, or ANOVA with post-hoc test correcting for false discovery (*alpha < 0.05, ** alpha < 0.01). Bar graph data are shown as mean +/− SEM.

**Fig. S3:**
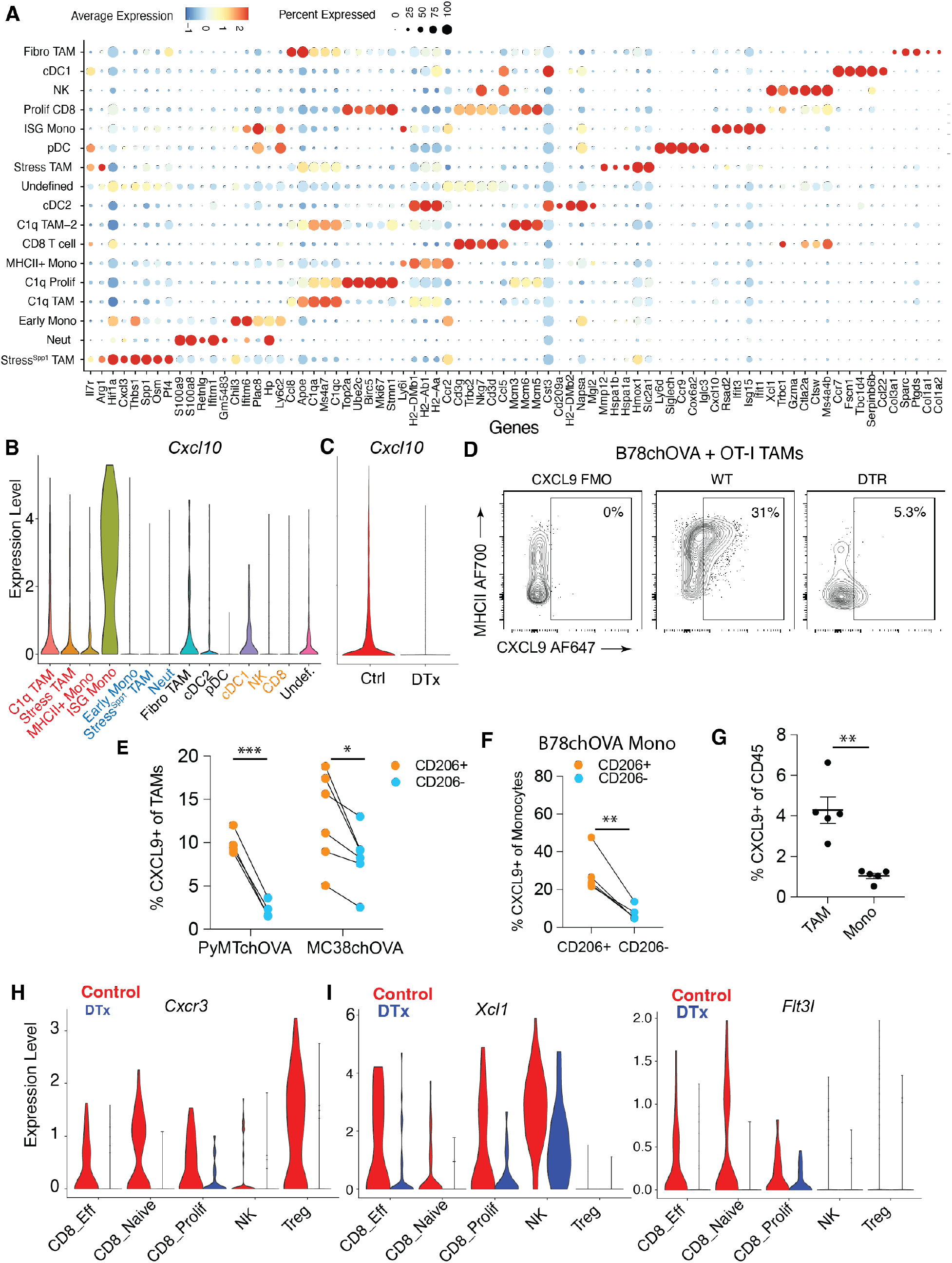
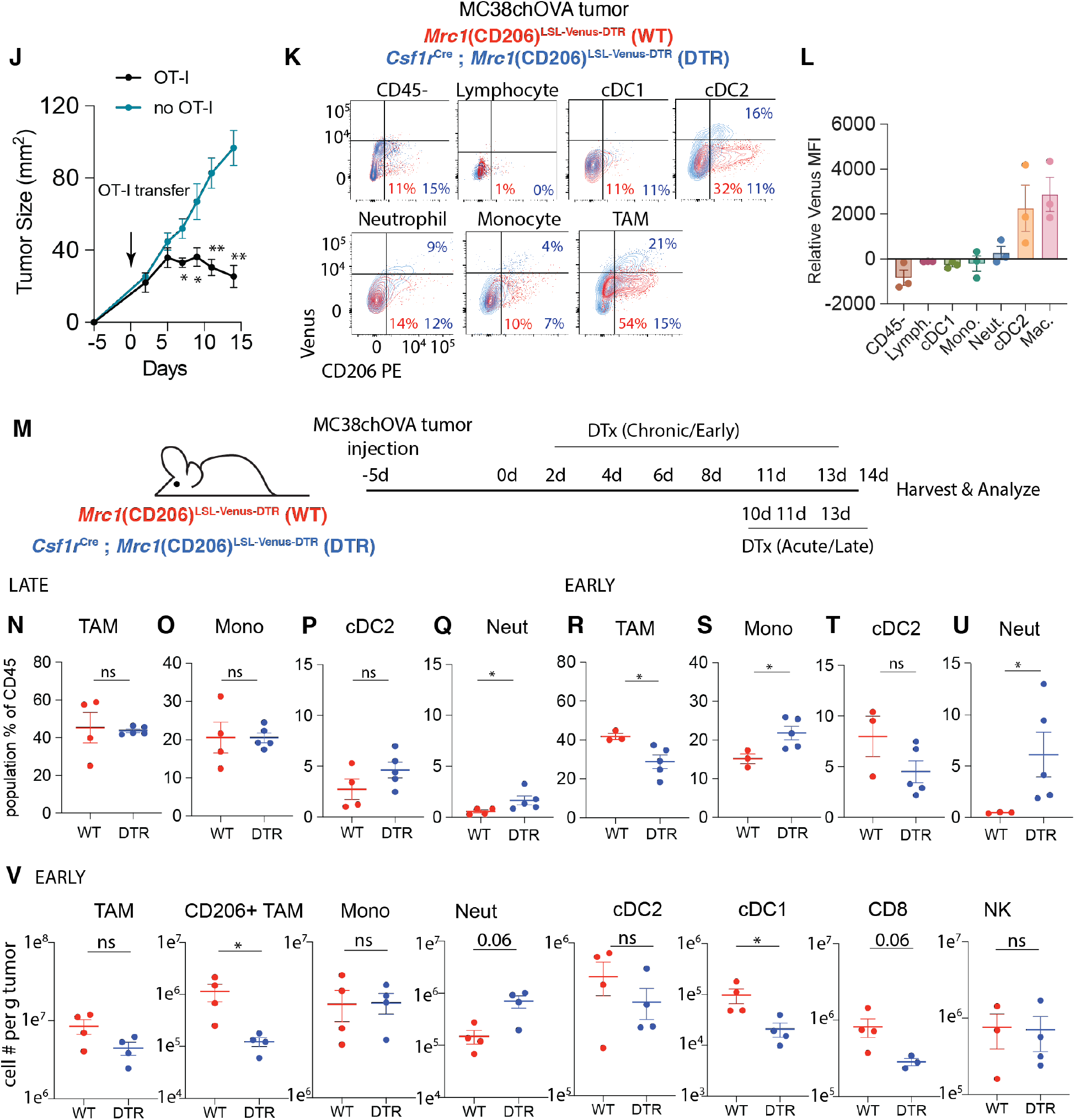
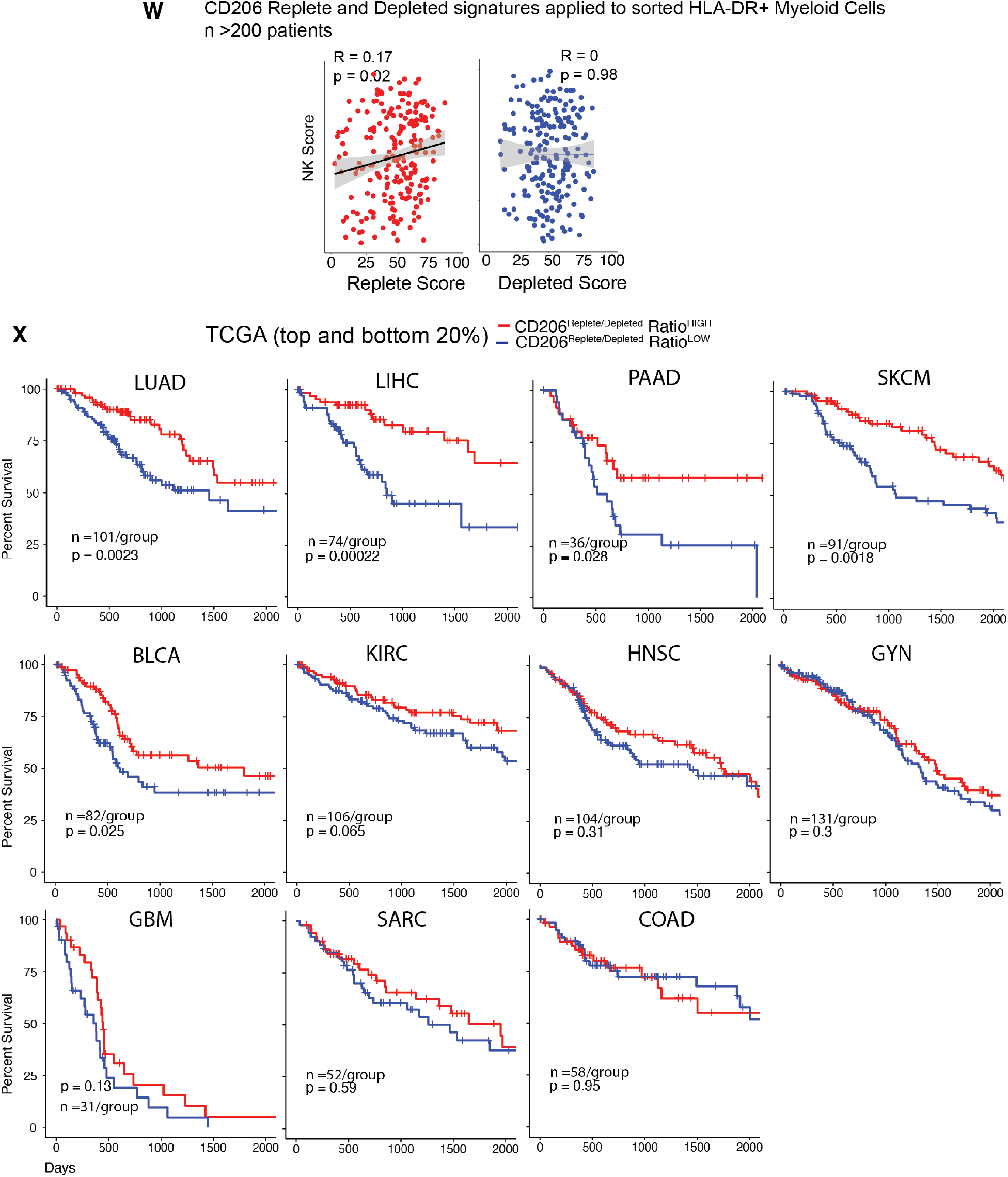
**(A)** Dotplot representing top5 differentially expressed genes and select other genes in each immune cell cluster identified from a harmonized dataset of spatially barcoded Control and DTx treated B78chOVA tumors d12 post adoptive transfer of CD2dsRed; OT-I cells; *Cxcl10* expression **(B)** aggregated across treatment conditions by cluster and **(C)** aggregated across clusters by treatment; **(D)** Representative flow cytometry plots showing CXCL9 expression in B78chOVA TAMs with or without DTx mediated depletion; **(E)** CXCL9 expression in PyMTchOVA and MC38chOVA (both without OT-I adoptive transfer) TAMs split by their CD206 expression; **(F)** CXCL9 expression in OT-I treated B78chOVA (d14 post adoptive transfer) monocytes and **(G)** relative abundance of CXCL9+ TAMs and Monocytes in the same context; Violin plot representing **(H)** *Cxcr3*, **(I)** *Xcl1* and *Flt3l* expression in the lymphoid compartment in Control and DTx treated conditions; **(J)** Representative time course of MC38chOVA tumor size with or without adoptive transfer of OT-I T cells; **(K)** Overlaid flow cytometry plots showing reporter (Venus) and CD206 expression in different immune cells in MC38chOVA tumors in WT (red; *Mrc1*^LSL-Venus-DTR^) and DTR (blue; *Csf1r*^Cre^; *Mrc1*^LSL-Venus-DTR^) mice and **(L)** quantification of relative reporter expression (DTR – WT) in the different subsets. **(M)** Schematic representation of the experimental setup for early and late CD206+ TAM depletion in MC38chOVA tumors using *Mrc1*^LSL-Venus-DTR^ (WT) and *Csf1r*^Cre^; *Mrc1*^LSL-Venus-DTR^ (DTR) mice; Relative abundance of different immune populations as a percentage of CD45+ cells with **(N-Q)** late and **(R-U)** early depletion regimens. **(V)** Abundance of different immune populations as total number of cells per g of MC38chOVA tumor in WT and DTR mice by the early DTx administration regimen; **(W)** Scatter plots of the CD206^Replete^ and CD206^Depleted^ Mono/Mac score per patient with the NK cell score (Pearson R and p value for the null hypothesis that there is not a correlation are noted); **(X)** Kaplan-Meier survival curves of patients grouped by the value of the CD206^Replete^: CD206^Depleted^ signature ratio (top and bottom 20%) from TCGA split by indications, p values for the log-rank test are noted for each curve in (X) bar graph data are mean +/− SEM, ****p<0.0001, **p<0.01, *p <0.05, ns = no significance by paired ratio t-tests (E, F) or unpaired t-tests or Mann-Whitney test.

## Notes

### Competing Interest Statement

The authors have declared no competing interest.

### Summary of Updates

Added new data regarding: -Compensatory neutrophils do not account for the immune suppressive effect of CD206+ TAM depletion -Detailed analysis of CXCL9 expression in CD206+ vs. CD206- TAMs and Monocytes in tumors; -Ability of CD206+ vs. CD206- TAMs to differentially attract CD8 T cells -Additional Fig. 5: detailed delineation of CD206Replete and Depleted signatures and prognostic relevance to human cancer indications -Supplementary data reorganized into 3 figures

